# Decades of Trivers-Willard research on humans: what conclusions can be drawn?

**DOI:** 10.1101/2022.08.22.504743

**Authors:** Valentin Thouzeau, Jeanne Bollée, Alejandrina Cristia, Coralie Chevallier

**Affiliations:** LNC^2^, Département d’études cognitives, École normale supérieure, Université PSL, INSERM, 75005 Paris, France; LSCP, Département d’études cognitives, École normale supérieure, Université PSL, EHESS, CNRS, 75005 Paris, France

**Keywords:** Systematic review, Trivers-Willard Hypothesis, Human Evolution, Sex-ratio, Investment

## Abstract

The Trivers-Willard hypothesis predicts that parents in good condition are positively biased towards sons, while parents in poor condition are positively biased towards daughters. An extensive literature testing this hypothesis has accumulated in the last five decades. We take stock of results concerning humans in a systematic review, which yielded 87 articles, reporting a total of 821 hypothesis tests. A p-curving analysis did not reveal a pattern of p-values consistent with p-hacking. Effects are consistent with the Trivers-Willard hypothesis overall. We then went on to check whether there was a difference between sex ratio and post-birth investment. Theoretical work suggests that, the conditions under which the Trivers-Willard hypothesis is verified should be more restrictive in the case of post-birth investment than for sex ratio. We explored this question in two ways and obtained mixed results. We put forward recommendations for future studies that aim to further assess the validity of the Trivers-Willard hypothesis or mechanisms subtending it, and we discuss the implications of different ways of measuring parental status and investment.

## 1. Introduction

The Trivers-Willard hypothesis is a classic in evolutionary biology (Salmon & Hehman, 2020; Trivers & Willard, 1973). The theory initially put forward by Robert L. Trivers and Dan E. Willard predicts that parents in good condition will be positively biased to invest more in male offspring, whereas parents in poor condition will be positively biased to invest more in female offspring. These predictions follow from the fact that (a) the reproductive success of offspring depends on parental conditions, and (b) the reproductive success of males is more variable than that of females. Under these premises, parents in good conditions have an adaptive advantage to invest in their sons since they are more likely to have many offspring than daughters, whereas parents in poor conditions have an adaptive advantage to invest in their daughters since they are more likely to guarantee offspring than sons^1^. More than 4,000 papers have cited this hypothesis since the original paper was published almost 50 years ago. Based on this large body of research a number of advances and remaining pitfalls can be highlighted.

First, additional theoretical work has clarified the exact conditions under which the predictions should hold. In particular, what is meant by “parental condition” and “investment” was not entirely clear in the original paper. Recent work has since emphasized that investment could take two forms, (1) biased sex ratios and (2) biased post-birth parental care towards one or the other sex, with evidence that the former should be seen in a wider range of settings than the latter (Veller et al., 2016).^2^

Second, several systematic reviews have assessed the explanatory power of the Trivers-Willard hypothesis by formally integrating the results of the literature in non-human species. Overall, these studies provide mixed results and it remains unclear whether the Trivers-Willard hypothesis applies to all species. Hewison and Gaillard (Hewison & Gaillard, 1999) for instance, surveyed studies on ungulates, finding mixed results, which provided weak but significant support for the hypothesis in a subsequent meta-analysis (Sheldon & West, (Sheldon & West, 2004). In 2001, Brown focused on parental investment in non-human primates, but concluded that it was difficult to accurately test the hypothesis because adequate measures of offspring’s increased reproductive success through maternal care were lacking in these species. Similarly, Cameron (Cameron, 2004) and Douhard (Douhard, 2017) reviewed studies on sex ratio in non-human mammals, and reported that findings were highly variable across species. Finally, narrative reviews suggest that the results are also mixed among humans (Cronk, 2007).

This overall picture of mixed results is unsurprising given that the effect predicted by Trivers and Willard is very small, which mechanically increases the probability of false negatives. In this context, systematic review approaches are particularly useful because they leverage data aggregation across studies, including those reporting null effects, and can be less biased than narrative reviews (Bishop, 2020; Thomas-Odenthal et al., 2020). Many fields of scientific research have also noted that null results are under-reported (Bakker & Wicherts, 2011; Loken & Gelman, 2017), which means that even systematic reviews can wind up capturing a mere publication bias. Such a bias can reflect scientists’ pressure to selectively publish positive findings (Rosenthal, 1979) or researchers’ bias to selectively report analyses if the resulting p-value falls below the usual 5% significance level (Simonsohn et al., 2014b), a bias known as *p-hacking* (Head et al., 2015).

This paper builds on the extensive literature testing the Trivers-Willard hypothesis in humans through a systematic review that avoids selection bias (PRISMA, 2020), combined with p-curving to check for the possible presence of p-hacking (Simonsohn et al., 2014a). Considering the small effect predicted by the Trivers-Willard hypothesis, and the large variety of tests and methodologies, it is difficult to extract the data necessary to calculate effect sizes (see Table S3). We therefore focused on analyses based on p-values. Although these do not allow the estimation of effect sizes, they are informative in terms of the presence of p-hacking bias as well as the consistency of effects. Our analyses take into account the conceptually diverse operationalizations of (a) parental condition and (b) parental bias (sex ratio and parental investment). Note that operationalizing parental condition and investment taking into account construct validity (Grahek et al., 2021) is key to testing the hypothesis empirically (Cronk, 2007). Therefore, even though theories only clearly distinguish sex ratio from post-birth investment, we will suggest additional categorizations for both parental condition and parental bias below.

## 2. Methods

### 2.1. Literature screening and coding

#### 2.1.1. Inclusion criteria

We considered articles written in English and published in peer-reviewed journals. Since the Trivers-Willard hypothesis has a very identifiable name, we restricted our bibliographical research to that phrase. After keyword exploration, we made queries in Google Scholar (November 6th 2020, incognito mode; after 200 results, only a negligible number of articles actually tested the Trivers-Willard hypothesis, so we stopped the screening after 300 results following current recommendations (Haddaway et al., 2015)) with the keywords “Trivers-Willard + human + children + mean + test”, and in PubMed with the keywords “Trivers-Willard-animal” (October 9th 2020). We also performed a backward search by adding the articles cited explicitly as testing the Trivers-Willard hypothesis in humans in previously selected articles.

#### 2.1.2. Exclusion criteria

During the *Screening phase*, one author (JB) manually excluded articles whose abstract made it clear that the Trivers-Willard hypothesis was not tested, as well as other inclusion criteria noted above (articles that were not in English, or not peer-reviewed). Next, we searched the entire document for the keyword “Trivers-Willard”, automatically or visually depending on the format of the file. We excluded studies whose abstract and occurrences of Trivers-Willard in the main text showed that the article did not explicitly test the Trivers-Willard hypothesis on humans (i.e., tested animals, hypothesis not mentioned or not tested, review or theoretical papers).

During the *Eligibility phase*, we excluded articles that did not provide a direct test of the Trivers-Willard hypothesis as such, e.g., articles focusing on proximal mechanisms (thus assuming that the hypothesis is true), testing an extension of the hypothesis (see (Anonymous, 2022) for more details), or bringing up the hypothesis only in the discussion (so that the connection appears post hoc). We also excluded papers that reported no usable p-values, that were a review, or that reported over 200 tests (which would have unduly affected results).

During the *Analyses phase*, we conducted a p-curving study and an exploratory study. In the p-curving study, we excluded articles that did not report at least one exact p-value and that reported only non-significant p-values. We also chose to exclude significant tests going against the predictions of the hypothesis. Indeed, if the Trivers-Willard hypothesis is supported by p-hacking evidence, then results that run counter to the hypothesis cannot be the result of p-hacking aimed at supporting the classical hypothesis. In the exploratory study, we also excluded articles reporting only test results that were contrary to the predictions of the hypothesis.

#### 2.1.3. Data extraction

The first and second authors independently extracted the data from five randomly sampled articles (Mackey, 1993; Mealey & Mackey, 1990; Sparks, 2016; Valente, 2015; Wallner et al., 2012), and checked that the instructions led to identical extractions. Then, the first author extracted the information of the remaining articles.

For each selected article, we collected: title, date of publication, author list, and journal. We then extracted every statistical test of the Trivers-Willard hypothesis within each article. Whenever a test was not clearly related to the prediction, it was not included. For example, if there was a list or table of p-values, and at the end the authors concluded with “overall, our results support the Trivers-Willard hypothesis”, those p-values could not be included in the database because it was unclear which ones were relevant according to the coders.

For each statistical test, we collected the country of origin of the sample, the sample size, the sampling unit (mothers, fathers, children, or families), the data collection procedure (e.g., interviews, anthropometric measurements, demographic surveys), the measure of parental condition (e.g., mother’s age, family SES; see exhaustive list in Table S1 in (Anonymous, 2022)), the measure of investment (e.g., duration of breastfeeding, birth weight, or inheritance; see exhaustive list in Table S2 in (Anonymous, 2022)), and whether the measure of investment was drawn from parents (such as time fathers spend with children) or children (such as birth weight).

We also collected information specific to each statistical test: control variables, type of test (e.g., Chi-square, Probit regression, event history model; exhaustive list in SI), p-value (in the approximate form as “< 0.05” or the exact form as “0.0012”), and whether the result of the test was non-significant or significant (always using an uncorrected alpha = .05), and in the latter case whether results aligned with the Trivers-Willard hypothesis’ predictions or not. Where information was ambiguous (e.g., when p-values were reported without reference to a clearly identified test, or when the type of test was not reported) or missing, it was reported as missing data.

#### 2.1.4. Coding of categories for exploratory analyses

For exploratory purposes, we split the types of parental condition into five general categories (see full table of correspondences in SI): condition measured at the **macroscopic** scale (such as life expectancy in a country or whether a country has been exposed to a famine); **economic, educational and material resources** (such as income, SES or number of years of education); **social status** in a family or a group (such as marital status, domestic violence or belonging to a minority ethnicity); **biological condition** (such as body mass index, number of children or maternal age), and **experimentally-induced condition** (such as having been exposed to disturbing news in a laboratory experiment).

We also split post-birth investment into four general categories: biological measures of parents (such as duration of lactation), behavioral measures of parents (such as giving one’s wealth to an heir), biological measures of the child (such as birth weight), and measures of the child related to the assumed outcome of the investment (such as the educational attainment of the child).

## 3. Results

All statistical analyses and graphical representations were generated with the software *R* version 4.0.3 (R Core Team, 2013), and the packages *ggplot2, ggpubr, lmerTest* (scripts available in (Anonymous, 2022)).

Our search resulted in 84 articles that could be included in analyses, with inclusion and exclusion criteria as detailed in Figure 1. Although a majority of samples were based in North America, the geographic coverage was considerable, and data on diverse operationalizations of parental bias (sex ratio, investment or both) comes from a wide range of sites: Bangladesh, Canada, China, Cuba, Czech Republic, Dominica, England, Ethiopia, Gambia, Germany, Ghana, Hungary, Ifaluk, India, Japan, Kenya, Mexico, Namibia, Nepal, Netherlands, Papua New Guinea, Philippines, Poland, Rwanda, Slovakia, Spain, Sweden, Tanzania, Uganda, UK, USA, and Venezuela, in addition to 10 with data collected in several places, and 2 based on historical data.

**Figure 1.**
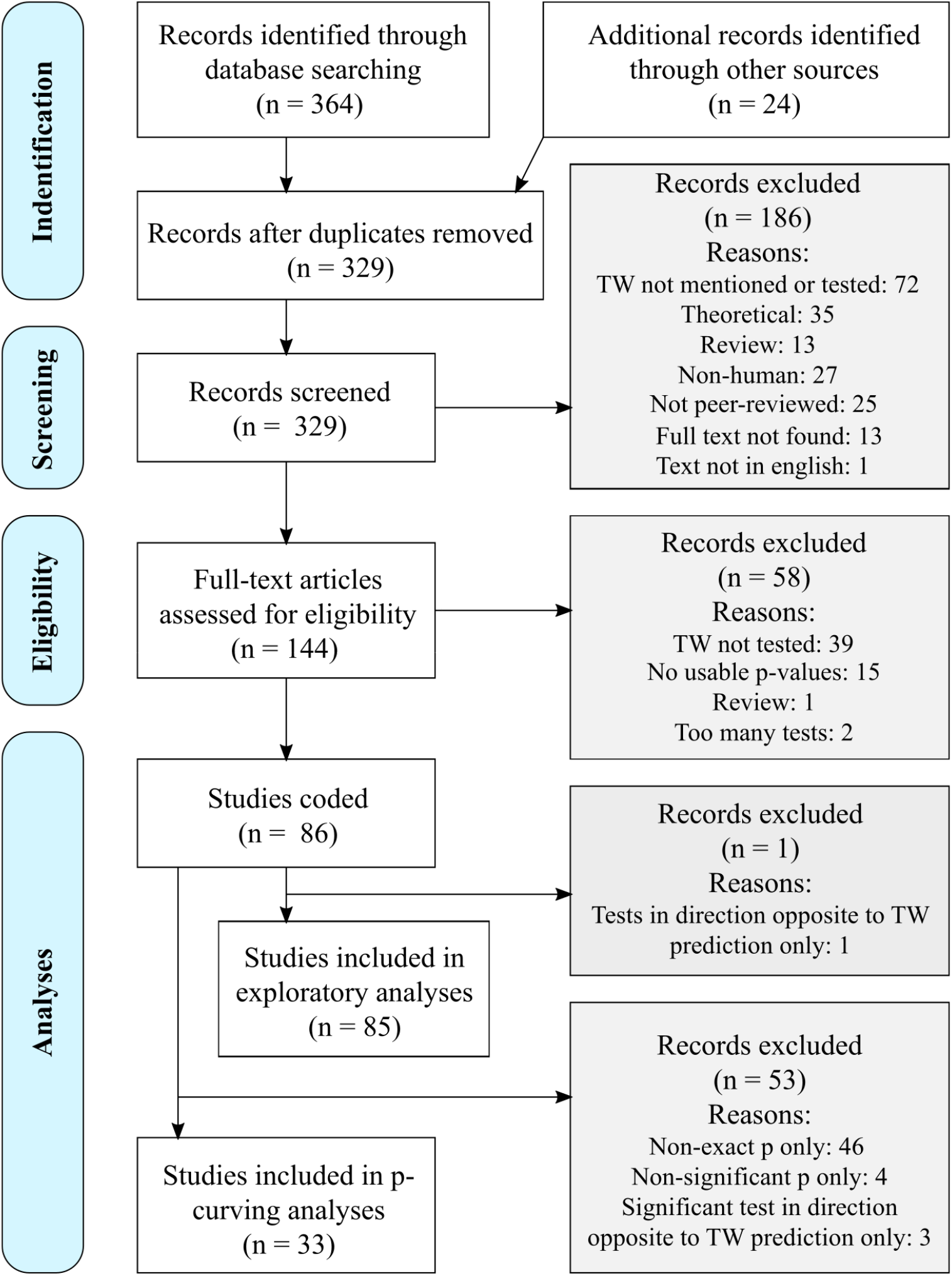
PRISMA flow chart describing database searches results, number of abstracts screened, number of full text screened, and exclusion criteria.

Figure 2 describes how the literature is distributed in temporal and disciplinary terms. The number of publications has increased in line with the overall increase in scientific publications, and with no clear association between year of publication and the operationalization of parental bias (sex ratio or investment). A majority of the included articles appeared in interdisciplinary, anthropology and biology journals, with a few articles appearing in journals classified as bearing mainly on demography, economics, psychology and sociology. We also observe a trend for an association between venue and parental bias measure: A majority of articles in anthropology, psychology, and sociology journals use parental post-birth investment measures, whereas sex ratio tests tend to be published in biology or interdisciplinary journals.

**Figure 2.**
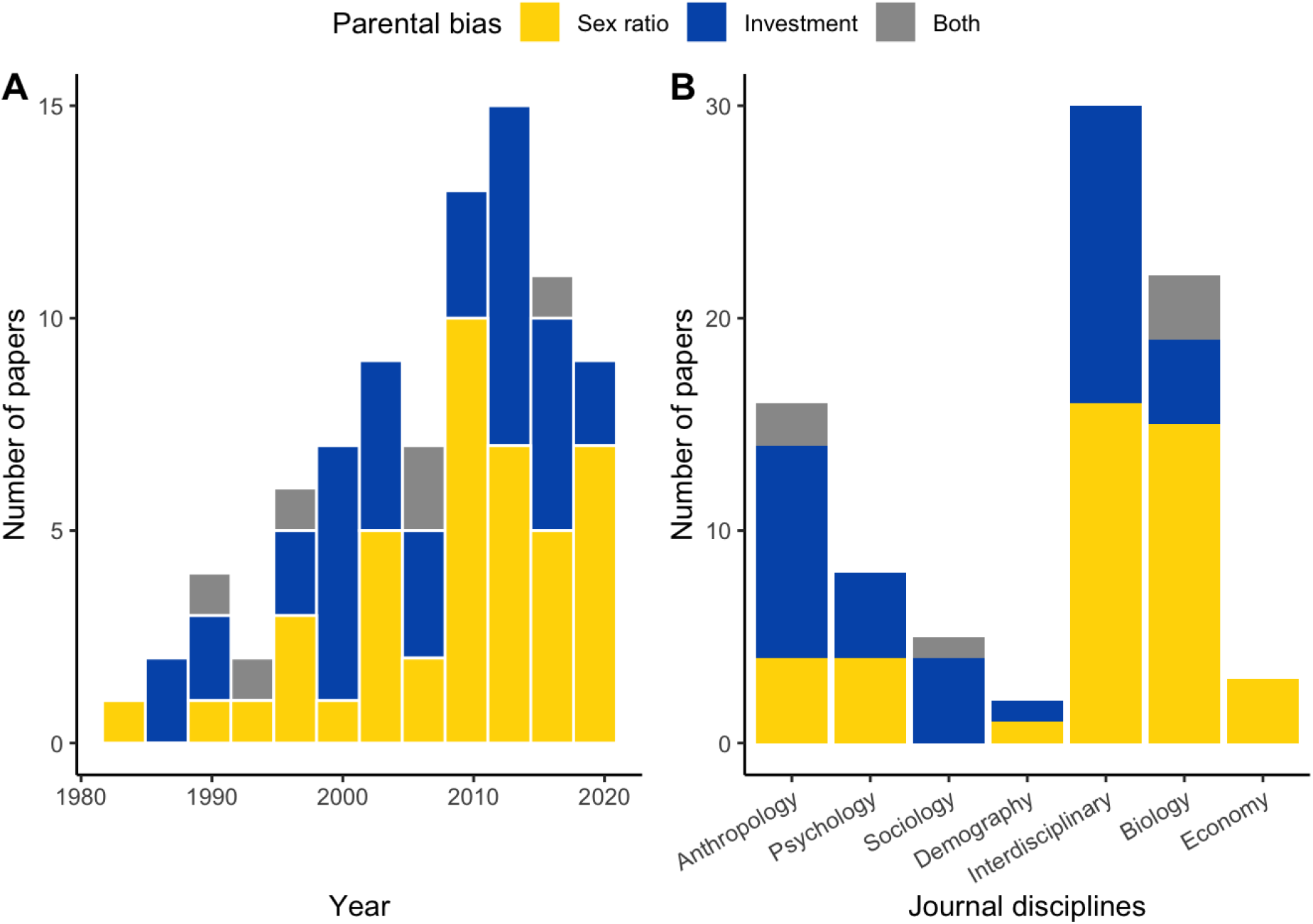
Number of papers as a function of type of parental bias and **A)** year of publication. **B)** journal discipline.

### 3.1. Checking for the presence of p-hacking

We first address the main question of whether there is evidence for the Trivers-Willard hypothesis and/or reporting biases. This first analysis builds on p-curving methods (Simonsohn et al., 2014b), which measure p-hacking in the literature and the robustness of an effect, by looking at the distribution of significant p-values between 0 and 0.05 (the typical alpha level) at 0.01 increments (i.e., 0.01, 0.02, 0.03, 0.04 and 0.05). This *p-curve* is expected to be flat if the effect does not exist, and if there is no p-hacking. The p-curve is expected to be strictly decreasing (i.e. the majority of significant p-values are closer to 0) if there is an effect. If there are many values at the 0.05 boundary, then the p-curve is consistent with p-hacking. The intuition behind this is that authors may (albeit inadvertently) select analyses pipelines leading to results that are significant at the 0.05 level. Following recommended procedures, we did not include approximate p-values in this analysis (of the form “< 0.01” for example). This resulted in a total of N = 94 p-values. We then randomly drew one p-value per article in order to have independent samples, and repeated this procedure 100 times before plotting the distribution of the p-curves in Figure 4.

**Figure 3.**
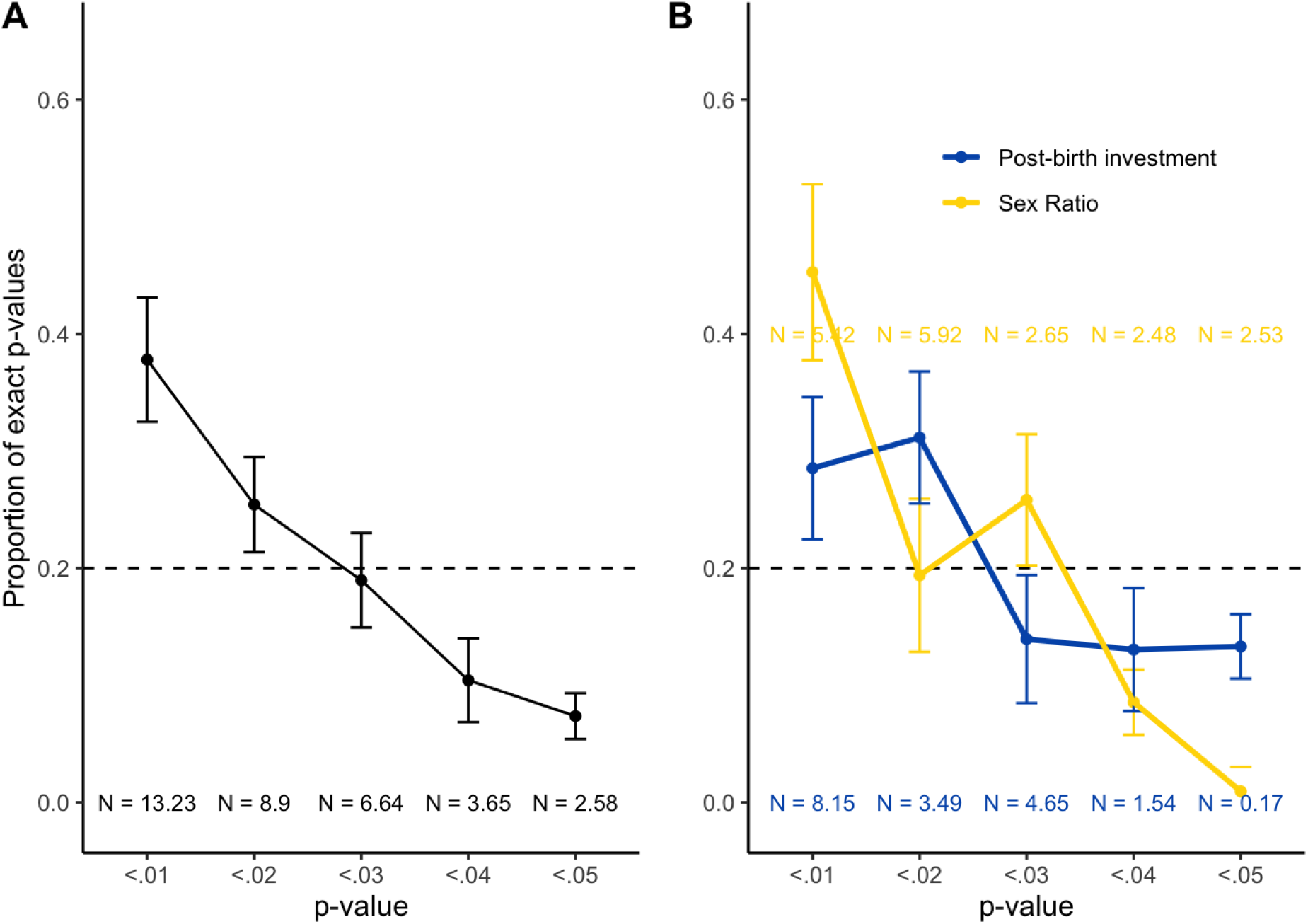
Average p-curves over 100 replicates describing the distribution of the exact p-values lower than 0.05 by type of study, for **A.** all p-values (black), and **B.** separate types of p-values by sex ratio (blue) and post-birth investment (orange). The horizontal dotted line indicates what is expected in the case of a null effect without p-hacking. The error bars indicate the standard deviation from the 100 random draws of p-curves. The Ns indicate the average number (across 100 replicates) of p-values in each significance category.

**Figure 4.**
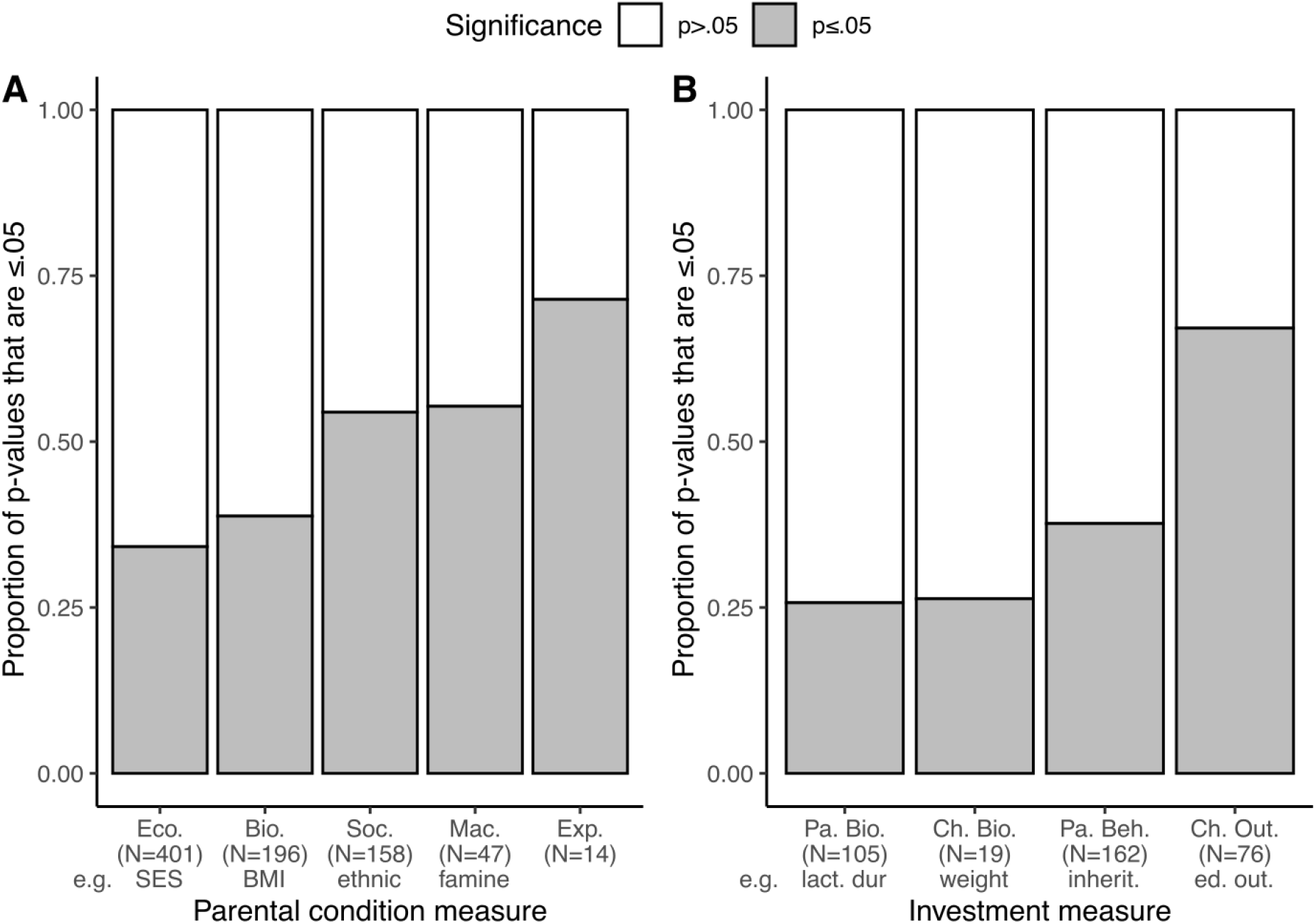
Proportion of p-values that are significant as a function of **A)** the different ways of measuring parental condition (**Eco.** = Economic and educational condition, e.g., maternal SES; **Bio**. = Biological condition, e.g., BMI, body mass index. **Soc** = Social status related condition, e.g., belonging to an ethnic minority; **Mac**. = Macroscopically measured conditions, e.g., country hit by famine; **Exp**. = Experimentally induced condition, e.g., receiving disturbing news) and **B)** the different ways of measuring investment (**Pa. Bio.** = biological measures of parents, e.g., lactation duration; **Ch. Bio.** = biological measure of the child, e.g., birth weight; **Pa. Beh.** = behavioral measure of parents, e.g., wealth inherited to the child; **Ch. Out.** = measure of the child related to the outcome of the investment, e.g., child’s educational outcomes).

Figure 3 thus shows traditional p-curves, using only the exact p-values grouped in 0.01 increments. Overall, results are most consistent with no p-hacking and with the presence of an effect.

Given theoretical arguments that there may be differences for the two operationalizations of parental bias, we separated two groups of p-values. Both curves seem right skewed, which is compatible with evidence supporting the Trivers-Willard hypothesis and not with reporting biases. Visually, the right skew of the sex ratio tests is more pronounced than that of the investment tests.

As the Trivers-Willard hypothesis assumes a different variance of reproductive success between males and females, it is possible that the observed results depend on marriage rules. Indeed, it has been shown that societies with monogamous marriage rules lead to a closer variance in reproductive success between males and females, in contrast to polygamous societies where reproductive success yields a very different variance between the sexes (Brown et al., 2009). One might therefore expect the hypothesis to be valid only in societies with polygamous marriage rules.

To assess this possibility, we listed the marriage rules of each society studied in the included articles, referring to the legal norms of each country. We then replicated our analyses by eliminating all studies conducted on populations from countries where polygamy is legally permitted. The results obtained only on the legally monogamous populations alone were qualitatively identical to the previous results (see Figure S1). This indicates that marriage rules do not determine the effect assumed by the Trivers-Willard hypothesis. This is probably caused by the fact that, although some human populations have monogamous marriage rules, actual behaviors do not necessarily align with those prescribed by the rules.

### 3.2. Exploratory analyses of subtypes of parental condition measures and investment measures

Figure 4 shows the frequency with which different types of measures were used, and highlights the variety of ways in which parental condition and investment are operationalized. The most common measures of parental condition were those associated with economic and educational status, followed by biological condition and those related to social status. Only a few studies measured parental condition on a macroscopic scale (i.e. region-wide, like a country-wide famine) or induced them experimentally. To better inform future work, we performed exploratory analyses to dig deeper into our database and test whether there is evidence mostly consistent with the Trivers-Willard hypothesis regardless of the study’s methodology.

The analyses described above only build on significant p-values, which is not all of the tests that have been provided in the literature, and thus do not give us enough power to address this important question. Since our analyses above did not reveal any sign of p-hacking, we took a different tack here, by including all p-values reported in all studies and considering both significant (N = 336) and non-significant (N = 486) p-values.

To check whether the operationalization of parental condition and parental investment mattered, we constructed a generalized binomial mixed model to predict whether a test was significant or not (if p ≤ .05 or p > .05), from the type of parental investment (sex ratio or post-birth investment) and the type of parental condition as fixed variables, and the study identifier as a random variable. All 797 p-values from 84 papers could be included in this analysis. Parental bias (sex ratio or investment) did not predict the proportion of significant tests (ß = −0.04, SE = 0.39, p = 0.91), whereas parental condition did. Specifically, economic and educational condition of the parents, taken as baseline for the test, was associated with the lowest proportion of significant tests (34%, N = 401). Biological measures of condition led to non-significantly higher proportions (39%, N = 196, ß = 0.45, SE = 0.3, p = 0.14). The economic and educational baseline was significantly lower than that of tests where parental condition had been measured from social status (54%, N = 158, ß = 1.07, SE = 0.31, p = 0) and macroscopically (54%, N = 48, ß = 1.39, SE = 0.68, p = 0.04). There was no significant difference between parental condition established through economic and educational means and studies using experimental induction of condition (71%, N = 14, ß = 0.88, SE = 1.58, p = 0.58), but this null result may be rather due to high variance in the latter and low power in this comparison because of the small number of p-values from studies using experimental induction.

A second regression predicted whether a test was significant or not among studies on investment specifically (351 p-values from 41 papers), now declaring as fixed effects the precise operationalization of post-birth investment. Studies measuring post-birth investment biologically on the parents actually resulted in the lowest proportion of significant outcomes (26%, N = 105), and were therefore declared as the baseline for this analysis. Against this baseline, children’s biological measures led to a non-significantly higher percentage (26%, N = 19, ß = −0.96, SE = 0.72, p = 0.18). Differences with the baseline of parental biological measures were significant for parents’ behavioral measures (37%, N = 163, ß = 0.85, SE = 0.58, p = 0.14), as was that for studies using children’s economic or educational status as the index for parental investment (67%, N = 76, ß = 1.97, SE = 0.73, p = 0.01).

### 3.3. Effect size estimation

To complement the p-curving analysis, we sought to estimate the effect size of the bias predicted by the Trivers-Willard hypothesis. The type of statistic performed as well as the way in which these can be combined through meta-analyses depend on whether it concerns investment or sex ratio. In the case of investment, studies never report an investment ratio (e.g., investment in boys divided by investment in girls), but instead report on the results of an interaction test (i.e., investment in boys versus girls as a function of parental condition). Estimating the size of this type of effect faces a series of obstacles (Simonsohn et al., 2022). While research developing effect size metrics is abundant for main effects, it is scarcer for interaction terms: Effect sizes are well-studied for e.g., a two-by-two matrix using chi-square tests, but a lot less for other tests (see e.g., Borenstein, 2009). Additionally, experts criticize the use of transformations to include studies using diverse statistical analyses in the same meta-analysis (Lakens, 2013). We found there was too much heterogeneity and too few results to hope that results of a meta-analysis on investment may be useful.

We thus turned to sex ratio, where there was a small but not negligible group of studies employing two-factor chi-square tests. We computed the effect size from 33 tests from 10 different studies using the *chies* function of the R package *compute.es*, and integrated them via a random-effects meta-regression using the R function *robu* from the *robumeta* package, which allows consideration of correlated structures (due to several effects coming from the same study). Our model accounts for variation in individual studies for effect size estimation through random effects. The overall effect size estimate was r = 0.037, (CI 95% = [0.0055; 0.0676], t = 2.84, p = 0.0275). A graphical evaluation of the funnel plot indicates a strong correlation between the effect and the sample size: the larger the sample, the smaller the effect (see Figure S2 in Anonymous, 2022).

## 4. Discussion

Our systematic review of the literature revealed that the Trivers-Willard hypothesis has spread outside evolutionary biology, and is now widely studied in various disciplines ranging from anthropology and sociology to demographics and economics, with samples from a variety of populations around the world. Our review found that tests on sex ratio and post-birth investment are similarly widespread, although tests on sex ratio are more frequent in biology journals and post-birth investment studies in anthropology and sociology journals.

Importantly, a p-curving analysis revealed no evidence of p-hacking. Further analyses showed that the effects predicted by the Trivers-Willard hypothesis were supported by the literature as a whole (Figure 3). We then checked whether there was a difference between sex ratio and post-birth investment. Theoretical work suggests that the conditions under which the Trivers-Willard hypothesis is verified should be more restrictive for post-birth investment than for sex ratio (Veller et al., 2016). We explored this question in two ways and obtained mixed results. On the one hand, we found that the right skew of the p-curve is visually more pronounced for sex ratio than it is for investment. On the other hand, an exploratory regression revealed that there was a similar proportion of significant effects for sex ratio and post-birth investment measures.

We also ran exploratory analyses to test whether different measures of parental condition and parental bias produced different results. Parental condition and parental investment have been operationalized in many different ways but it is not clear that all these measures have equal construct validity and are equally relevant to test the Trivers-Willard hypothesis. For instance, some measures may be influenced by uncontrolled factors, some may not predict ultimate reproductive success or affect boys and girls equally. Regarding parental condition, we found that measures of parents’ individual economic and educational status resulted in significant tests the least. By contrast, measures related to the status of a social group (e.g., being a single woman, or belonging to a dominant ethnic group), and even more so measures taken at a macroscopic scale (e.g., pre-versus post-famine), were more likely to support the Trivers-Willard hypothesis. This gradient suggests that more general measures of parental condition validate the hypothesis more often than measures of condition taken at the individual level. Regarding post-birth investment, biological measures (particularly those drawn from the child) led to the lowest proportion of significant p-values, whereas child outcome measures (e.g., economic or educational status attained by the child) led to the highest.

Taken together, these results suggest that support for the Trivers-Willard hypothesis is better detected with global measures than with narrower measures. This may indicate that, although the Trivers-Willard hypothesis is valid, it does not reflect a simple effect resulting from a single mechanism, but it emerges instead from a variety of causal pathways, whose very small individual effects need to accumulate to be statistically detectable. If this interpretation is correct, research on mechanisms will require markedly higher precision and power than what previous work allows.

We also estimated the size of the effect predicted by the Trivers-Willard hypothesis in the case of sex ratio. The effect, as one would expect, is particularly small (r = 0.037). A graphical evaluation of the funnel plot indicates a correlation between the effect and the sample size: the larger the sample, the smaller the effect. This could be related to a selection bias in the literature, but the p-curving analysis shows that the literature is not particularly p-hacked. This pattern may also be caused by the fact that researchers have a choice between: (1) studying a very large sample size with demographic data, but without accurate and relevant measures of parental conditions, or (2) focusing on a small sample size, which allows them to collect accurate and relevant measures of parental conditions. Under these conditions, it is expected that the effect size will be larger for studies with fewer participants, even without underlying p-hacking. Thus, this particular result does not contradict our p-curving analysis as a whole, which suggests no strong evidence for p-hacking once sample sizes are set aside and the whole literature is integrated.

Our meta-regression is limited in that a very small proportion of the literature could be included, but we think it may be informative to bear in mind this estimated size of effect for potential future work. We find that the confidence interval of the estimated effect is larger for studies with small sample sizes, and smaller for studies with large sample sizes. This is expected and similar to what is observed by Brown and Silk (2002) for non-human primates, because smaller samples are typically less precise. Taking all studies together, giving greater weight to larger and more precise studies, allows us to calculate an overall confidence interval of [0.0055; 0.0676], which leads us to reject the hypothesis that the effect is non-significant across studies.

Our results have a range of implications for future research on the Trivers-Willard hypothesis in humans. First, it appears that there is robust support for predictions for sex ratio, which calls for further work on proximal mechanisms among humans (Douhard, 2017). Sex ratio modulation occurs near conception in ungulates (Sheldon & West, 2004), and in several other non-human mammals (Cameron, 2004). In line with the cellular biology work of Larson et. al. (Larson et al., 2001), Cameron proposed that increased glucose levels would be responsible for inhibiting female blastocyst development. It has been shown that female bovines with higher testosterone levels have a higher proportion of male offspring (Grant & Irwin, 2005), as testosterone may signal dominance in the group. It has also been shown that glucocorticoid production in females in the human species in response to stress leads to greater embryonic mortality of male fetuses (Navara, 2010). These studies provide some interesting leads that need to be substantiated before we can reach a consensus on the proximal mechanisms of the sex ratio bias in the human species.

Second, our results may be interpreted as lending weaker statistical support for the Trivers-Willard hypothesis in terms of post-birth investment. We hypothesize that part of the problem here is one of construct validity. For example, it is not clear that maternal milk is a form of parental investment that ultimately affects the child’s reproductive success, or whether choosing to feed one’s infant poorly is a form of post-birth abortion that would fall more on the side of sex ratio manipulation. Documenting the whole causal chain will require multi-decade studies testing the relationship between parental condition, parental biological and behavioral measures, and the ultimate impact on children’s reproductive success. Although such work is difficult and expensive, large-scale longitudinal studies will be very important in the future (Berg et al., 2014; Nettle, 2002), and we trust that our exploratory results can contribute to designing them more efficiently. We therefore believe that large-scale studies would benefit from collecting data specifically on these variables. Moreover, these measures are linked to features on which public policies can act and could pave the way to more effective and more targeted interventions.

Before concluding, we address two potential limitations of our study. First, the empirical distinction between studies related to the sex ratio and those related to investment is generally not problematic, but some cases are ambiguous. In particular, the study of parental behaviors that are likely to increase or decrease infant mortality can be conceived either as a way of modulating the sex ratio, or as the by-product of a differential investment between boys and girls. We chose to consider such measures taken at or before birth as belonging to the sex ratio category, and those taken after birth as belonging to the investment category. Other distinctions may be equally relevant, however. Because the majority of cases are unambiguous, we believe that the conclusions we draw from our study are robust. Second, it is possible to categorize differently the types of parental condition and types of investment, and in particular to propose more numerous and thus more refined categories. We chose to use broad categories in order to limit ambiguity in the categorization process, to produce an analysis of the literature that is as intelligible as possible, and to ensure adequate statistical power. Future work could further investigate differents types of parental condition and investment in order to test which proximal phenomena validate Trivers-Willard hypothesis. In order to allow collaborative work (as proposed by (Tsuji et al., 2014)), we archived our database, so that the scientific community can update it as new evidence is produced, and propose alternative categorisations according to their interests. This deposit may be useful for future attempts at estimating effect sizes, although as obvious in Supplementary Information material, a meta-analytic approach may be impossible given that standardized effect sizes are not definable for a majority of the analyses (Borenstein et al., 2021). Third, the initial Trivers-Willard hypothesis has recently been complemented by theoretical work that clarifies what can be expected according to the ecology of the populations (Schindler et al., 2015). In particular, some conditions may lead to predictions contrary to the initial Trivers-Willard hypothesis. We chose to test whether the predictions of the hypothesis initially formulated and tested in the literature are validated in the case of the human species. It would be particularly relevant to carry out future work to measure all the ecological conditions that could lead to alternative predictions.

To conclude, our study is the first to exhaustively assess the validity of the Trivers-Willard hypothesis, proposed almost 50 years ago, with a focus on the human species. We provide clear evidence that this literature can be trusted, with no sign of p-hacking. Further analyses suggest that there might be interesting modulations as a function of important conceptual and empirical distinctions. These modulations will need to be substantiated by longitudinal studies, which are now needed to test the mechanisms that are at play in the Trivers-Willard effect.

## 5. Contributions

JB performed the inclusion phase.

VT performed the backward phase.

VT and JB elaborated the coding scheme and performed the initial data extractions to verify the reproducibility of the instructions.

VT performed the data extraction and coding into categories.

VT& AC performed the statistical analyses and produced the tables and the figures.

VT, JB, CC and AC wrote the paper.

CC & AC supervised the work.

## 6. Acknowledgments

We thank Laudine Carbuccia for her help in validating the categorization criteria and for helpful discussions. We thank the support of FrontCog ANR-17-EURE-0017 for its participation in funding this research. AC acknowledges the support of the J. S. McDonnell Foundation Understanding Human Cognition Scholar Award; European Research Council (ERC) under the European Union’s Horizon 2020 research and innovation programme (ExELang, Grant agreement No. 101001095). CC acknowledges the support of the ANR (ANR-21-CE28-0009).

## 7. Additional informations

The authors declare that they have no conflict of interest. All data and code are available at (Anonymous, 2022).

## Footnotes

1 The Trivers-Willard hypothesis is construed at an ultimate level: it assumes that a range of mechanisms (whatever they may be) regulate sex ratio and/or investment according to parental conditions. The proximal way in which these mechanisms are implemented, e.g., through sex-dependent abortions under conditions of limited food availability (Post et al., 1999), is not specified by the hypothesis.

2 The sex ratio is expected to be modified by parental condition whenever this condition is correlated with the reproductive success of the offspring. By contrast, investment is expected to change only under more restrictive conditions, namely that the reproductive success gain from investing a unit of parental care is more profitable for one sex than the other. As a result, theoretical studies predict that the Trivers-Willard hypothesis might apply for sex ratios but not for post-birth parental investment when the latter condition is not met.

